# Deoxyhypusine synthase is required for the translational regulation of pancreatic beta cell maturation

**DOI:** 10.1101/2023.04.24.537996

**Authors:** Craig T. Connors, Emily K. Anderson-Baucum, Spencer Rosario, Catharina B.P. Villaca, Caleb D. Rutan, Paul J. Childress, Leah R. Padgett, Morgan A. Robertson, Teresa L. Mastracci

**Affiliations:** Department of Biology, Indiana University Purdue University Indianapolis, Indianapolis, IN 46202, USA; Indiana Biosciences Research Institute, Indianapolis, IN 46202, USA; Department of Biostatistics and Bioinformatics, Roswell Park Comprehensive Cancer Center, Buffalo, NY 14203, USA; Department of Biochemistry and Molecular Biology, Indiana University School of Medicine, Indianapolis, IN 46202, USA; Center for Diabetes and Metabolic Diseases, Indiana University School of Medicine, Indianapolis, IN 46202, USA

**Keywords:** beta cell maturation, mRNA translation, translational regulation, beta cell development, beta cell identity, beta cell function, diabetes

## Abstract

As professional secretory cells, beta cells require adaptable mRNA translation to facilitate a rapid synthesis of proteins, including insulin, in response to changing metabolic cues. Specialized mRNA translation programs are essential drivers of cellular development and differentiation. However, in the pancreatic beta cell, the majority of factors identified to promote growth and development function primarily at the level of transcription. Therefore, despite its importance, the regulatory role of mRNA translation in the formation and maintenance of functional beta cells is not well defined. In this study, we have identified a translational regulatory mechanism in the beta cell driven by the specialized mRNA translation factor, eukaryotic initiation factor 5A (eIF5A), which facilitates beta cell maturation. The mRNA translation function of eIF5A is only active when it is post-translationally modified (“hypusinated”) by the enzyme deoxyhypusine synthase (DHPS). We have discovered that the absence of beta cell DHPS in mice reduces the synthesis of proteins critical to beta cell identity and function at the stage of beta cell maturation, leading to a rapid and reproducible onset of diabetes. Therefore, our work has revealed a gatekeeper of specialized mRNA translation that permits the beta cell, a metabolically responsive secretory cell, to maintain the integrity of protein synthesis necessary during times of induced or increased demand.

**ARTICLE HIGHLIGHTS:**

- Pancreatic beta cells are professional secretory cells that require adaptable mRNA translation for the rapid, inducible synthesis of proteins, including insulin, in response to changing metabolic cues. Our previous work in the exocrine pancreas showed that development and function of the acinar cells, which are also professional secretory cells, is regulated at the level of mRNA translation by a specialized mRNA translation factor, eIF5A^HYP^. We hypothesized that this translational regulation, which can be a response to stress such as changes in growth or metabolism, may also occur in beta cells.
- Given that the mRNA translation function of eIF5A is only active when the factor is post-translationally modified (“hypusinated”) by the enzyme deoxyhypusine synthase (DHPS), we asked the question: does DHPS/eIF5A^HYP^ regulate the formation and maintenance of functional beta cells?
- We discovered that in the absence of beta cell DHPS in mice, eIF5A is not hypusinated (activated), which leads to a reduction in the synthesis of critical beta cell proteins that interrupts pathways critical for identity and function. This translational regulation occurs at weaning age, which is a stage of cellular stress and maturation for the beta cell. Therefore without DHPS/eIF5A^HYP^, beta cells do not mature and mice progress to hyperglycemia and diabetes.
- Our findings suggest that secretory cells have a mechanism to regulate mRNA translation during times of cellular stress. Our work also implies that driving an increase in mRNA translation in the beta cell might overcome or possibly reverse the beta cell defects that contribute to early dysfunction and the progression to diabetes.

## INTRODUCTION

As professional secretory cells, pancreatic beta cells require high-level, adaptable mRNA translation to facilitate the rapid synthesis of proteins, including insulin, in response to changing metabolic cues (1,2). Specialized mRNA translation programs are essential drivers of cellular development and differentiation (3,4). However, in the pancreatic beta cell, the majority of factors identified to promote growth and development primarily function at the level of transcription (5). Therefore, despite its importance, the regulatory role of mRNA translation in the formation and maintenance of functional beta cells is not well defined. The creation of regenerative therapeutics to prevent or treat diabetes, a disease characterized by loss or dysfunction of the insulin-producing beta cells, necessitates a comprehensive understanding of all mechanisms that drive functional beta cell development (6).

The endogenous generation of functional beta cells involves a multi-step and progressive developmental program. Many transcription factors, including Pdx1, Nkx6.1, Nkx2.2 and Neurod1, are essential for specifying beta cell fate during embryonic pancreas development (7). Additional maturation is then acquired in the postnatal period following the upregulation of several markers required for maintenance of cellular identity and function, including Slc2a2 (Glut2), MafA, and Ucn3 (8–11). The proteins needed for beta cell functional development are acquired in a timed manner that follows with the progression of the animal through expansion of beta cell mass in the perinatal period and acquisition of secretory machinery after weaning (12). Clearly, this well-regulated timeline requires coordination between transcriptional and translational machinery. However, the mechanisms regulated by mRNA translation are not well understood.

An important regulator of mRNA translation is eukaryotic initiation factor 5A (eIF5A) (13,14). Genome-wide association studies (GWAS) have identified *Eif5a* in a type 1 diabetes (T1D) susceptibility locus in humans and NOD mice (15,16), and the eIF5A pathway was shown to be active in islets during T1D (17). These findings suggest that eIF5A is of great significance to the beta cell, and that this translation factor may function to establish and/or maintain proper beta cell health (18). For eIF5A to perform its function in mRNA translation, it must be post-translationally modified by the rate-limiting enzyme deoxyhypusine synthase (DHPS) in a process known as hypusine biosynthesis. DHPS catalyzes the addition of a 4-aminobutyl moiety from the polyamine spermidine on to the lysine at position 50 of eIF5A, which generates the modified amino acid hypusine (N-4-amino-2-hydroxybutyl(lysine)) and thus the hypusinated form of eIF5A (eIF5A^HYP^) (19). The eIF5A^HYP^ form is considered the “active” form, and it functions in the cytoplasm to facilitate mRNA translation initiation, elongation, and termination (20–25). Genetic deletion of *Dhps*, and thus loss of eIF5A^HYP^, specifically in the embryonic pancreas alters mRNA translation, with subsequent specific reductions in the synthesis of proteins critical for exocrine pancreas growth and function (26,27). Moreover, inducible loss of *Dhps* in the adult beta cell in the setting of high-fat diet was shown to impair compensatory beta cell proliferation (28). Therefore there is growing evidence that mRNA translation facilitated by DHPS/eIF5A^HYP^ is a critical regulatory mechanism to produce and maintain function in secretory cells in the pancreas.

Herein, we have generated a mouse model with a beta cell-specific deletion of *Dhps* from the onset of beta cell differentiation in the embryo to determine the role of hypusine biosynthesis in beta cell development and function. Our findings reveal that beta cell DHPS is dispensable for perinatal beta cell development; however, activation of the specialized translation factor eIF5A by DHPS plays a critical regulatory role in maintaining postnatal beta cell identity and function.

## MATERIALS AND METHODS

### Animal studies

Animal studies were approved by the Indiana University School of Medicine Institutional Animal Care and Use Committee (IACUC) and the Indiana University-Purdue University Indianapolis School of Science IACUC. Mice containing the *Dhps^LoxP^* allele (B6.Cg-Dhps^tm1.1Mirm/J^) (28) were mated with *Ins1-cre* mice (B6(Cg)-Ins1^tm1.1(cre)Thor/J^) (29) to generate mutants (*Dhps^LoxP/LoxP^*;*Ins1-cre*, denoted as DHPS^ΔBETA^) and littermate controls. Given that the *Ins1-cre* is a knock-in allele, all matings were arranged to produce only animals heterozygous for this allele. To provide a lineage trace and confirm cre-mediated recombination, the *R26R^Tomato^* reporter (B6.Cg-Gt(ROSA)26Sor^tm14(CAG-tdTomato)Hze/J^) (30) was bred into the mouse model (Supplemental Figure 1 A-F). All animals were maintained on a C57Bl/6 background and genotyped as previously described (28–30).

### Metabolic analyses

Animals were weaned at 3 weeks-of-age. From weaning until 6 weeks-of-age, body weight and blood glucose were measured weekly using a digital scale (Fisher) and AlphaTrak2 glucose monitor and test strips (ADW Diabetes), respectively. At 4 weeks-of-age, male and female animals were fasted for 5 hours and a glucose tolerance test (GTT) performed. For the GTT, animals were injected intraperitoneally with 2 g/kg glucose (0.25g/mL stock solution; Fisher) and blood glucose measured from tail blood at 0, 15, 30, 45, 60, 90, 120 minutes after injection. Statistical significance was determined using a one-way ANOVA (Prism9, GraphPad) (31).

One week-old animals were also evaluated for body weight and *ad lib* blood glucose measurements as per above. Blood was collected into EDTA coated tubes (Fisher) and processed by centrifugation (2000 rcf, 5 minutes). Separate cohorts of animals at 4, 4.5, and 6 weeks-of-age were evaluated for blood glucose levels following a 5-hour fast; blood was collected from tail vein (6 weeks-of-age) or by decapitation (1 and 4 weeks-of-age) into EDTA coated tubes (Fisher) and processed as above. Plasma samples from animals at 1, 4, and 6 weeks-of-age were analyzed by ELISA for insulin (Alpco) and glucagon content (Mercodia). Statistical significance was determined using a Student’s t-test (Prism9, GraphPad) (31).

### Immunofluorescence, immunohistochemistry, and morphometric analysis

Pancreas tissue at 1, 4, and 6 weeks-of-age was fixed in 4% paraformaldehyde (Acros Organics), cryo-preserved using 30% sucrose (Fisher), embedded in OCT (Fisher), and sectioned (8 μm) using a CM1950 Cryostat (Leica). For 1 week-old samples, every 10^th^ section was evaluated across the entire pancreas totaling 6 of 60 sections; for 4- and 6-week-old pancreata, every 10^th^ section was evaluated across the entire pancreas totaling 10 of 100 sections. Immunofluorescence was performed as previously described (32). Primary and secondary antibodies are listed in Supplemental Materials (Table S1). Whole section tile scan and individual islet images were acquired using a 710 confocal microscope (Zeiss) or a C2 confocal microscope (Nikon Corporation). All islets were evaluated from every section imaged. For morphometric analysis, insulin and glucagon area were measured across the entire tissue section and normalized to exocrine area as visualized by CarboxypeptidaseA. Zen Blue V2.3 software (Zeiss) was used to calculate endocrine and exocrine area. For cell counting analyses, islet z-stack images were evaluated using NIS Elements Software V5.30.01 (Nikon). Islet area, shape (circularity and length), and proximity to ducts were measured using QuPath software (https://qupath.github.io/) (33). Islet and duct structures were identified with the pixel classifier, and distance between structures was measured from the center of each islet to the nearest duct. Statistical significance was determined using a Student’s t-test (Prism9, GraphPad) (31).

Immunohistochemistry was also used for morphometric analysis of pancreas tissue from 4- and 6-week-old animals. Briefly, tissue sections were washed in 0.3% hydrogen peroxide (Fisher) in phosphate buffered saline (PBS) (Fisher). Tissue sections were then permeabilized in PBS with 0.1% TritonX (Fisher) and blocked with 5% donkey serum in PBS. Primary and secondary antibodies are listed in Supplemental Materials (Table S1). Tissue sections were developed in 3,3’-Diaminobenzidine solution (Fisher) for 10 minutes. Hematoxylin (Sigma) was used as a counterstain to visualize tissue architecture. All sections were rinsed in water and dehydrated in ethanol (Decon Labs), cleared in xylene (Fisher), and mounted using Permount (Fisher). Insulin or glucagon area was normalized to total pancreas area. Zen Blue V2.3 software (Zeiss) was used to calculate endocrine and exocrine. Statistical significance was determined using a Student’s t-test (Prism9, GraphPad) (31).

### Western blot analysis

Pancreatic islets were isolated from mice at 4 weeks-of-age (34) (see Supplemental Materials) and evaluated by western blot analysis as previously described (26). Primary and secondary antibodies are listed in Supplemental Materials (Table S1). After incubation with secondary antibodies, membranes were washed with 1% Tween (Fisher) in PBS, and then imaged and quantified by densitometry using Image Studio Software (LICOR Biosciences). Densitometric data are graphed as relative expression; statistical significance was determined using a Student’s t-test (Prism9, GraphPad).

### Quantitative mass spectrometry and proteomic data analysis

Pancreatic islets were isolated from mice at 4 weeks-of-age and subjected to proteomic analysis by mass spectrometry. Islets were collected from *Dhps^loxP/loxP^;Ins1-cre;R26R^Tomato^* and *Ins1-cre;R26R^Tomato^* mice. The number of islets per sample ranged from 116 to 193 islets. Sample preparation and mass spectrometry analysis was performed in collaboration with the Indiana University Center for Proteome Analysis at the Indiana University School of Medicine as previously published (26). Specific details for islet sample preparation and peptide measurement by mass spectrometry are in Supplemental Materials.

Data was exported as RAW files and analyzed in Proteome Discover™ 2.4 (ThermoFisher) with FASTA databases including uniprot *Mus musculus* reviewed and unreviewed sequences plus common contaminants. Quantification methods used isotopic impurity levels (Thermo Fisher). SEQUEST HT searches were conducted with a maximum number of two missed cleavages; precursor mass tolerance of 10 ppm; and fragment mass tolerance of 0.6 Da. Static modifications used for the search were carbamidomethylation on cysteine (C) residues, and TMT six-plex label on lysine (K) residues and the N-termini of peptides. Dynamic modifications used for the search were oxidation of methionines and acetylation of N-termini. Percolator False Discovery Rate was set to a strict setting of 0.01 and a relaxed setting of 0.05. Values from both unique and razor peptides were used for quantification. Peptides were normalized by total peptide amount with no scaling. Resulting grouped abundance values for each sample type, abundance ratio values, and respective *p*-values (ANOVA) from Proteome Discover^TM^ were exported to Microsoft Excel. Differentially expressed proteins (DEPs) were deemed significant with a |logFC|>1.5 and a *p*-value < 0.05. DEPs were then functionally enriched utilizing the STRING database (35) to assess relationships between proteins, generate a network, and enrich for pathways in which differential proteins appear (36). The Wikipathways (37) plug-in for Cytoscape (38) was used to integrate and visualize functionally enriched pathways of interest based on protein level expression. The analysis was carried out in the R environment [4.0.1] for statistical computing.

## RESULTS

### *Dhps* is dispensable for perinatal beta cell development

Our previous work defined a role for DHPS/eIF5A^HYP^ in exocrine pancreas development and function (26). However, the extreme exocrine pancreas loss and reduced postnatal survival of the DHPS^ΔPANC^ mouse model (26) precluded further examination of beta cell growth, maturation, and function. Therefore to determine the role of DHPS specifically in the beta cell, we generated a mouse model with a genetic deletion of *Dhps* in the insulin-expressing cells from the onset of beta cell differentiation in the embryo (*Dhps^LoxP/LoxP^*;*Ins1-cre;R26R^Tomato^*, denoted DHPS^ΔBETA^). DHPS^ΔBETA^ mutant animals were viable and western blot analysis of isolated islets confirmed knockdown of DHPS concomitant with a significant reduction in eIF5A^HYP^ (Supplemental Figure 1 G-I).

Morphometric analysis was performed on pancreas tissue collected from DHPS^ΔBETA^ animals and *Ins1-cre* controls during two significant stages of beta cell growth: (1) the first week of life (postnatal day (P) 7), which marks significant expansion of beta cell mass, and (2) weaning, which is characterized by beta cell functional maturation (12,39,40). Beta cell mass in 1 week-old DHPS^ΔBETA^ mice and *Ins1-cre* controls was evaluated by immunofluorescence. Insulin-expressing beta cells and glucagon-expressing alpha cells were visualized, with no observable difference in islet architecture, size, or number between DHPS^ΔBETA^ mice and *Ins1-cre* controls (Figure 1A,B; Supplemental Figure 2). Moreover, quantification of beta cell mass also showed no significant difference between DHPS^ΔBETA^ mice and *Ins1-cre* controls (Figure 1C). Body weight, *ad libidum* fed blood glucose and plasma insulin and glucagon levels were also not significantly different between DHPS^ΔBETA^ and controls (Figure 1D-G). Altogether these data identify that the absence of *Dhps* in the beta cell does not impact perinatal beta cell expansion, islet architecture, or whole-body metabolism, suggesting that *Dhps* is dispensable for early postnatal beta cell growth and function.

**Figure 1.**
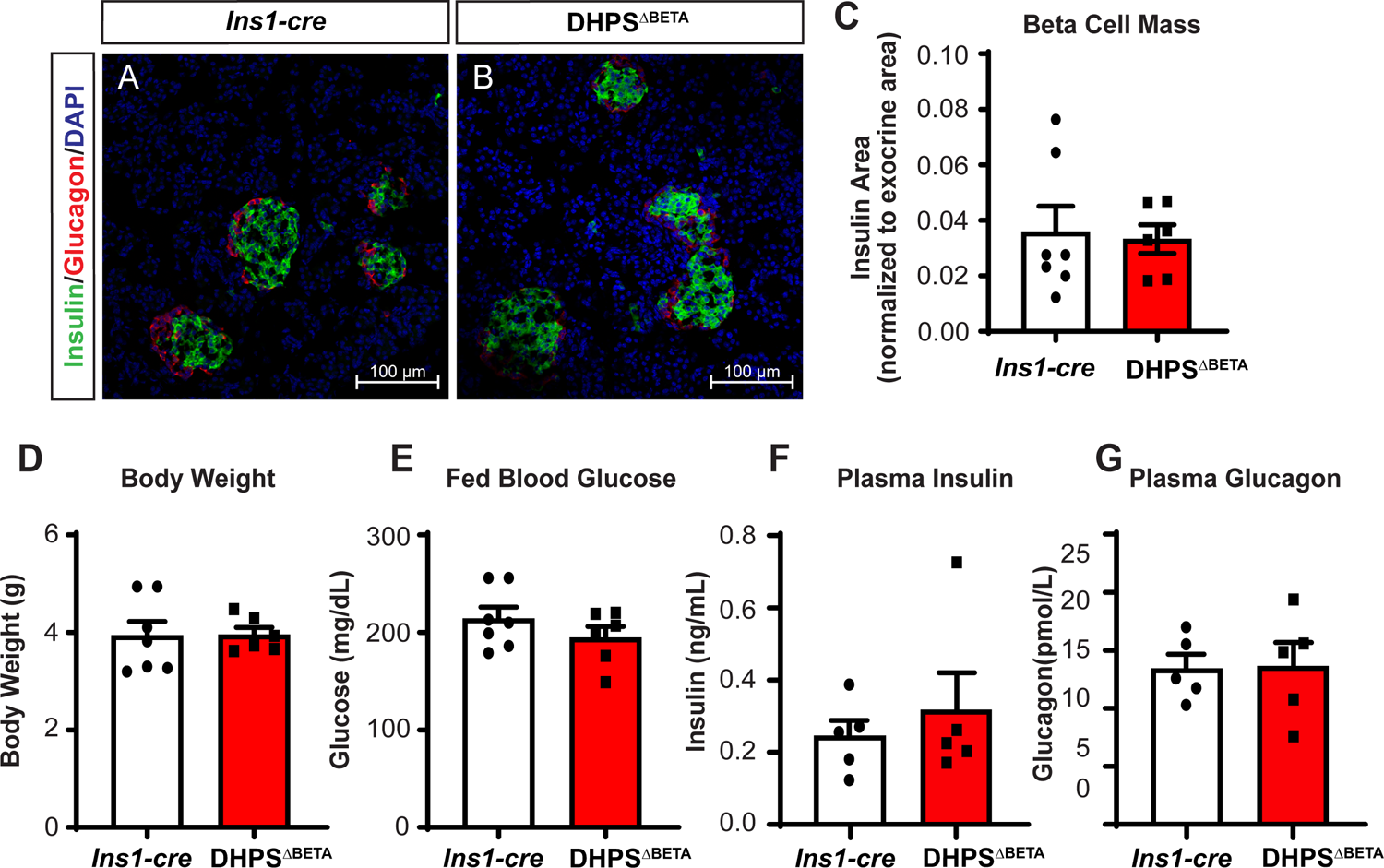
*Dhps* is dispensable for perinatal beta cell growth and function. Immunofluorescence for Insulin, Glucagon, and DAPI (nuclear marker) was performed on pancreas tissue from 1 week-old (A) *Ins1-cre* control and (B) DHPS^ΔBETA^ mutant mice. (C) Quantification of beta cell mass, normalized to exocrine area (n = 6 - 7/group). (D) Body weight and (E) *ad libidum* fed blood glucose at 1 week-of-age (n = 6 - 7/group). (F) Insulin and (G) Glucagon levels were measured from plasma collected from 1 week-old *Ins1-cre* and DHPS^ΔBETA^ mice (n = 5 - 7/group). Data are represented as mean +/- SEM.

### Deletion of beta cell *Dhps* results in the onset of diabetes after weaning

The second major stage of postnatal beta cell growth occurs at weaning (3 - 4 weeks-of-age). At this stage, animals experience a change in nutrition and their beta cells undergo a maturation that includes the acquisition of glucose-stimulated insulin secretion (12). Therefore we assessed DHPS^ΔBETA^ and *Ins1-cre* mice for changes in metabolic parameters at this stage of beta cell growth. Animals were evaluated beginning at 3 weeks-of-age, and we observed no significant difference in body weight between DHPS^ΔBETA^ and *Ins1-cre* animals until 6 weeks-of-age (Figure 2A). Whereas glucose tolerance tests at 4 weeks-of-age showed normal glucose response (Figure 2B,C), DHPS^ΔBETA^ mice dramatically progressed to hyperglycemia at 5 weeks-of-age and developed overt diabetes by 6 weeks-of-age (mean blood glucose of 675 mg/dL in males) (Figure 2D). Fasted blood glucose levels also confirmed the progression to diabetes, as DHPS^ΔBETA^ mice displayed elevated fasting blood glucose levels beginning at 4.5 weeks-of-age (mean blood glucose of 370 mg/dL in males), with an even greater defect observed at 6 weeks-of-age (mean blood glucose of 576 mg/dL in males) (Figure 2E). Male and female DHPS^ΔBETA^ and *Ins1-cre* mice were evaluated at all ages and identical phenotypes were observed in both sexes (see Supplemental Figures 3 and 4 for female data). These data identify that *Dhps* plays a critical role in postnatal beta cell function.

**Figure 2.**
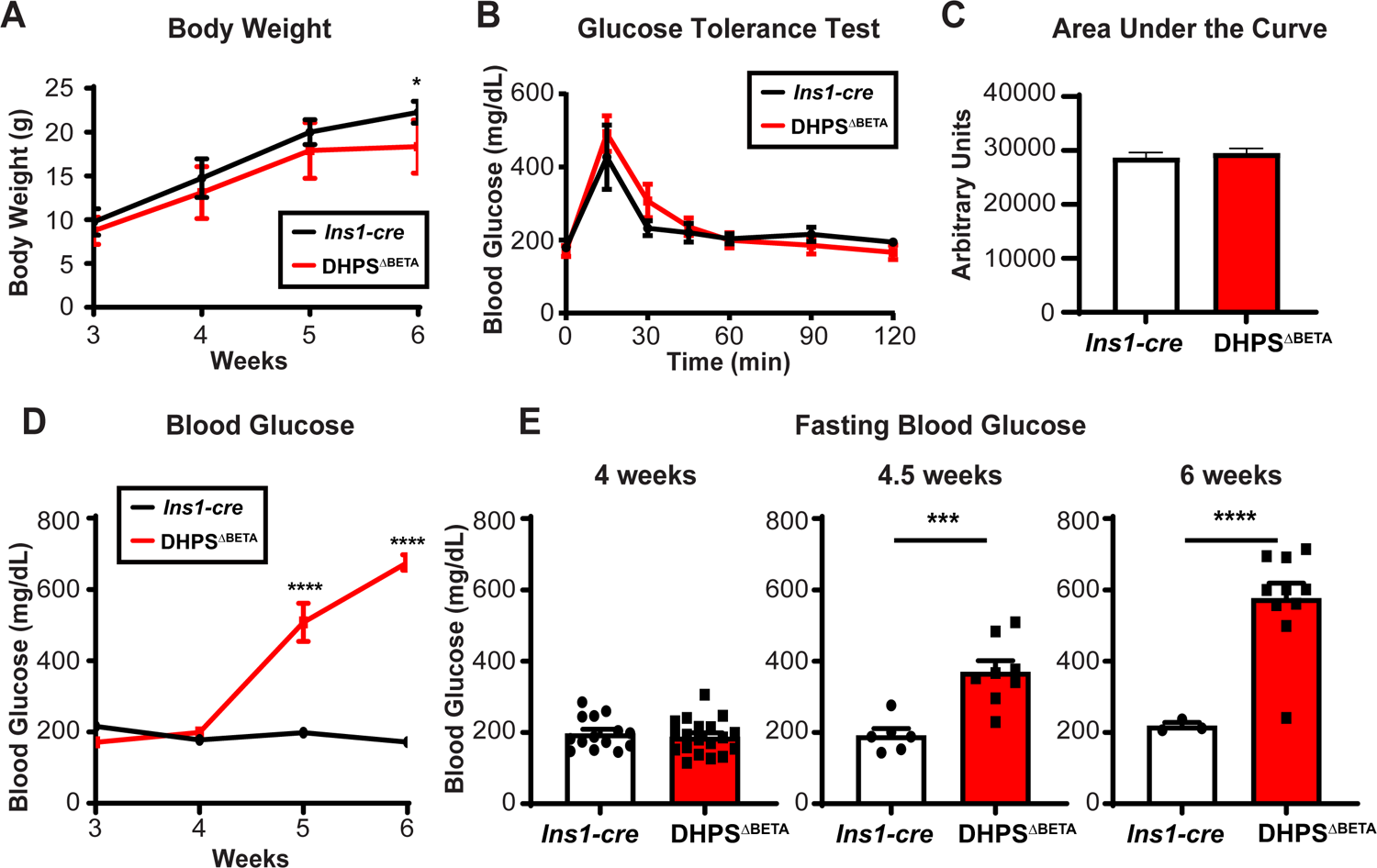
Measurement of metabolic parameters in the DHPS^Δ^^BETA^ mice after weaning. (A) Body weight was measured in *Ins1-cre* and DHPS^ΔBETA^ mice weekly beginning at weaning (3 weeks-of-age), (n = 4 - 8/group). (B) Glucose Tolerance Tests (GTT) were performed on 4-week-old mice (n = 4 - 8/group). (C) Area under the curve was quantified from the GTT data. (D) Blood glucose was measured weekly beginning at weaning (n = 4 - 8/group). (E) Blood glucose was measured after a 5-hour fast in 4-, 4.5-, and 6-week-old mice (n = 3 - 19/group). Data shown are for male mice only; data from female mice can be found in Supplemental Figure 3. Data are represented as mean +/- SEM. * p<0.05; *** p<0.001; **** p<0.0001.

### Beta cells in DHPS**^Δ^**^BETA^ mice have reduced insulin expression after 4 weeks-of-age

We showed that DHPS^ΔBETA^ mice develop hyperglycemia and diabetes after weaning. To determine the beta cell defect that contributes to this phenotype, we first performed morphometric analysis of pancreas tissue from DHPS^ΔBETA^ and *Ins1-cre* animals at 4 weeks-of-age, using immunohistochemistry to identify the insulin-expressing cells. Similar to what we observed at 1 week-of-age, islet structure at 4 weeks-of-age was not changed between the DHPS^ΔBETA^ mutants and *Ins1-cre* controls (Figure 3A,B) and quantification of beta cell mass showed no significant difference (Figure 3C). Moreover, blood collected from DHPS^ΔBETA^ and *Ins1-cre* mice at 4 weeks-of-age showed no difference in levels of plasma insulin or glucagon (Figure 3D, Supplemental Figure 4A). These results are in line with the normal glucose tolerance and fasted blood glucose levels at 4 weeks-of-age in the DHPS^ΔBETA^ mice (Figure 2B, C, E).

**Figure 3.**
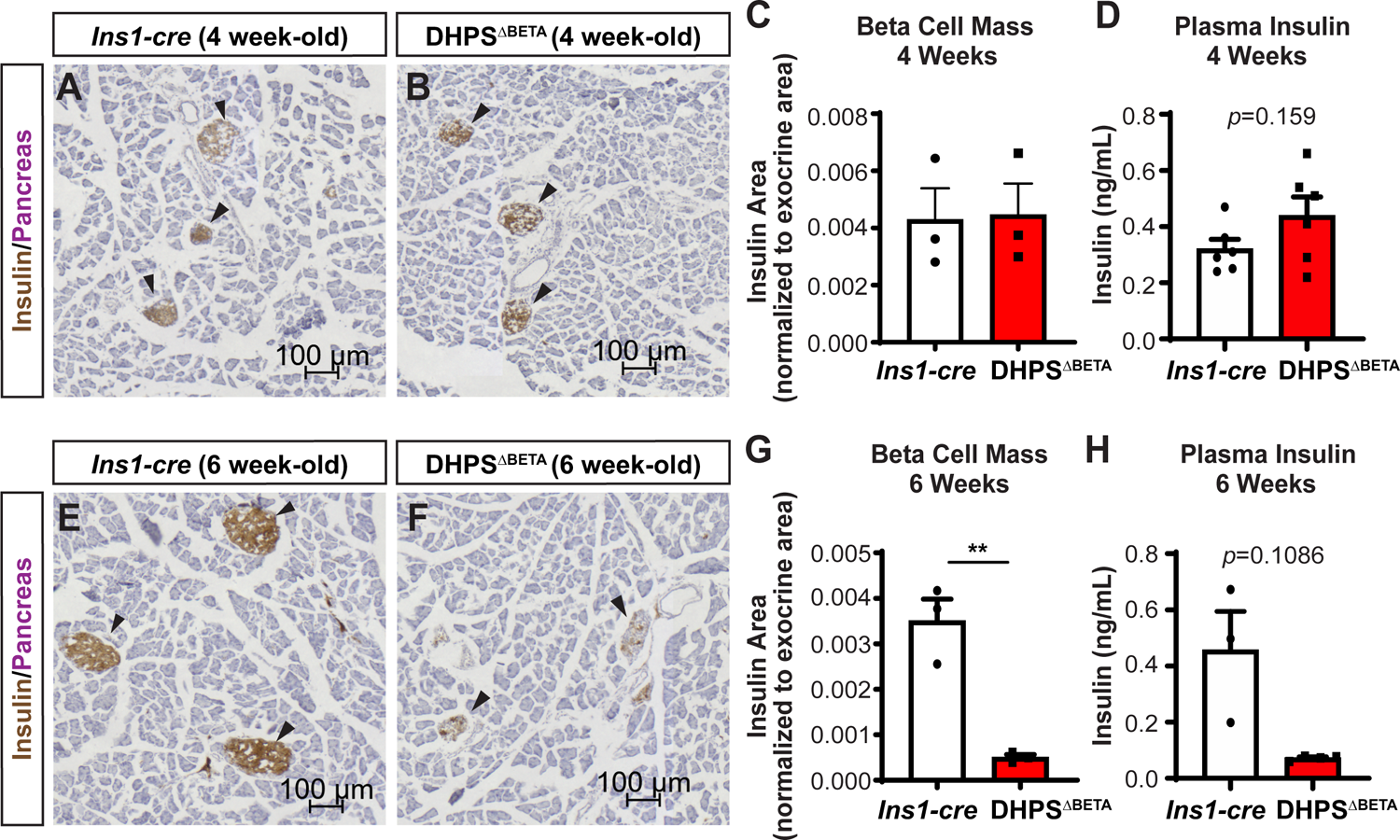
Beta cell mass in the DHPS^ΔBETA^ mice declines by 6 weeks-of-age Immunohistochemistry for insulin (brown) was performed on pancreas tissue from 4-week-old. (A) *Ins1-cre* control and (B) DHPS^ΔBETA^ mutant mice (n = 3/group). All sections were counterstained with Eosin (purple) to define pancreas tissue. (C) Beta cell mass was quantified as insulin area normalized to pancreas area. (D) Insulin levels were measured from plasma collected from 4-week-old *Ins1-cre* controls and DHPS^ΔBETA^ mutants (n = 6/group). Immunohistochemistry for insulin (brown) was performed on pancreas tissue from 6-week-old (E) *Ins1-cre* control and (F) DHPS^ΔBETA^ mutant mice (n = 3/group). All sections were counterstained with Eosin (purple) to define pancreas tissue. (G) Beta cell mass was quantified as insulin area normalized to pancreas area. (H) Insulin levels were measured from plasma collected from 6-week-old *Ins1-cre* controls and DHPS^ΔBETA^ mutants (n = 3-4/group). Data shown are for male mice only; data from female mice can be found in Supplemental Figure 4. Quantitative data are represented as mean +/- SEM. ** p<0.01

Because DHPS^ΔBETA^ mice were metabolically healthy at 4 weeks-of-age but rapidly demonstrated hyperglycemia and diabetes two weeks later, we examined pancreas tissue from DHPS^ΔBETA^ and *Ins1-cre* mice at 6 weeks-of-age. Using morphometric analysis, we identified a significant reduction in insulin-expressing cell mass in the DHPS^ΔBETA^ compared with *Ins1-cre* mice (Figure 3E-G). Concomitant with reduced beta cell mass, plasma insulin was reduced in DHPS^ΔBETA^ mice compared with *Ins1-cre* controls (Figure 3H). This change in beta cell mass and function did not impact the alpha cells, as the DHPS^ΔBETA^ mutants have normal alpha cell mass and no change in plasma glucagon, which was observed in both male and female mice (Supplemental Figure 4B-E). Altogether our data demonstrate that the absence of *Dhps* in the beta cell does not impact beta cell growth and function before 4 weeks-of-age; however, after this stage, significant beta cell dysfunction and diabetes occurs.

### Beta cells require DHPS to acquire cellular identity and maturation

DHPS is the rate-limiting enzyme required for the hypusination of eIF5A, which is the process by which eIF5A becomes post-translationally modified and thus active. The hypusinated/active form of eIF5A, eIF5A^HYP^, functions in specialized mRNA translation; however, the proteins that specifically require eIF5A^HYP^ for their synthesis in the beta cell have not been identified. We speculated that the phenotype of diabetes in DHPS^ΔBETA^ mice may be a result of altered specialized mRNA translation causing changes in the synthesis of certain critical beta cell proteins. Therefore, we used quantitative mass spectrometry on isolated islets from 4-week-old *Ins1-cre* controls and DHPS^ΔBETA^ mutants, which, importantly, is a stage when we observed no changes in beta cell mass or function. This analysis would determine, in an unbiased manner, the proteins whose synthesis is altered in the absence of *Dhps* in the beta cell. Overall, proteomic analysis detected 5900 proteins, with 123 significantly differentially expressed proteins (*p* < 0.05; |logFC|>1.5), both up- and down-regulated in the islets from DHPS^ΔBETA^ mice compared with *Ins1-cre* controls. Pathway analysis revealed alterations in numerous pathways critical for beta cell function including protein localization to the secretory granule, regulation of mRNA processing and regulation of cell cycle (Figure 4A). Moreover, detailed examination of this proteomic data showed a large number of critical beta cell proteins downregulated in the DHPS^ΔBETA^ islets (Figure 4B).

**Figure 4.**
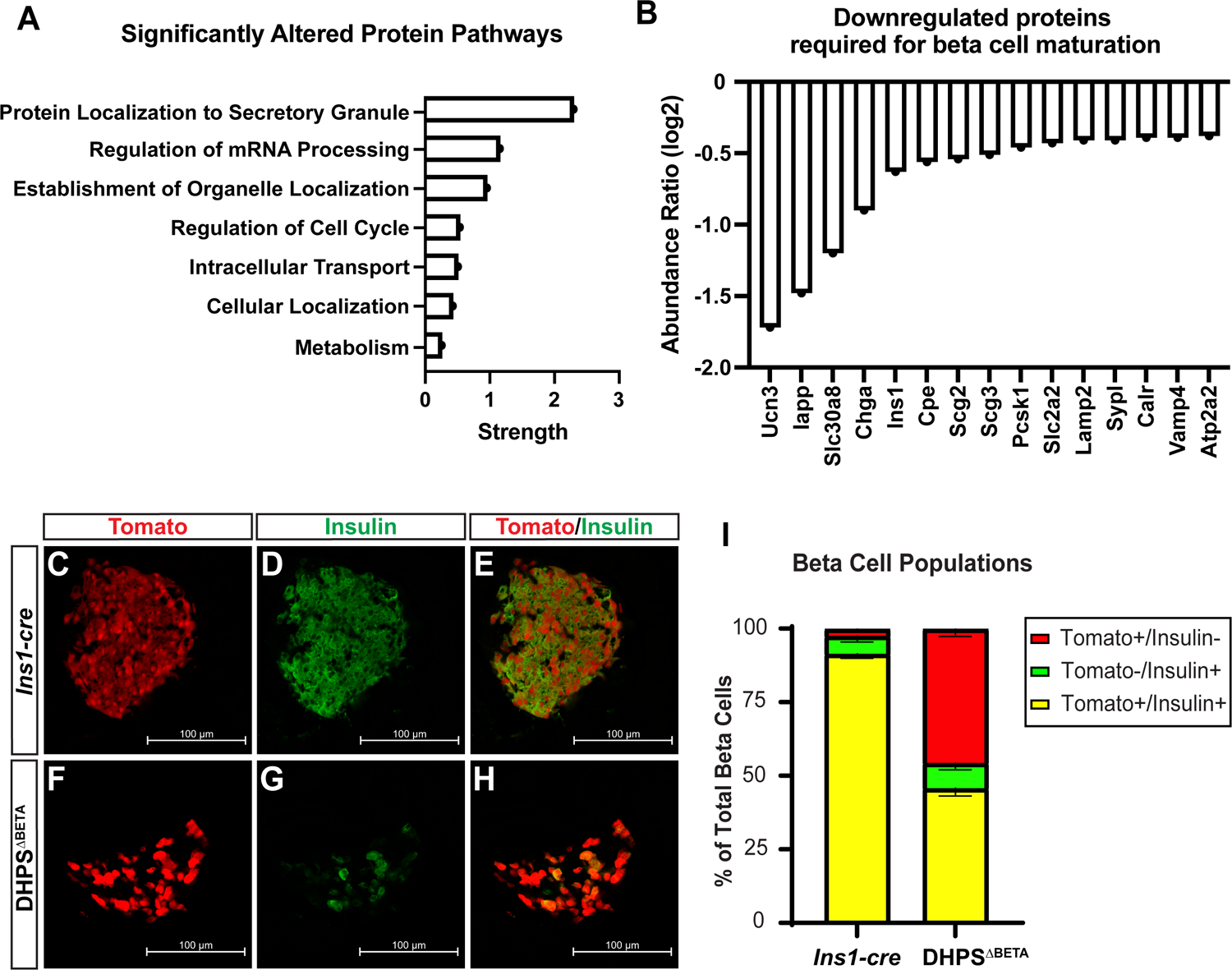
Islets from DHPS^ΔBETA^ mice show altered protein synthesis. (A) Pathway analysis performed on mass spectrometry data from 4-week-old DHPS^ΔBETA^ mutant islets compared with *Ins1-cre* controls. (B) Many proteins required for beta cell maturation were downregulated in the 4-week-old DHPS^ΔBETA^ mutant islets. Immunofluorescence was performed on pancreas tissue from 6-week-old (C,D,E) *Ins1-cre* and (F,G,H) DHPS^ΔBETA^ mice to determine the co-expression of insulin (green) with the *R26R^Tomato^*beta cell lineage reporter (Tomato, red) (n = 3/group). (I) Tomato+/Insulin-, Tomato-/Insulin+, and Tomato+/Insulin+ cell populations were quantified. Data are represented as mean +/- SEM.

Given that 4-week-old DHPS^ΔBETA^ mice are normoglycemic with unaltered beta cell mass, the differentially expressed proteins identified in these islets suggested that the reduction in beta cell mass observed 2 weeks later at 6 weeks-of-age could be the result of altered beta cell identity. Therefore, we used the *R26R^Tomato^* reporter allele to lineage-label insulin-expressing cells in DHPS^ΔBETA^ and *Ins1-cre* mice and analyze the beta cell characteristics in the islets at 6 weeks-of-age, which was when we observed the diabetes phenotype. As expected, wholemount imaging showed Tomato-expressing cells in the DHPS^ΔBETA^ mutant and *Ins1-cre* control isolated islet preparations, confirming the presence of cells that derived from the insulin-expressing cell lineage in both groups (Supplemental Figure 5). Morphometric analysis of pancreas tissue from 6-week-old DHPS^ΔBETA^ mutant and *Ins1-cre* control animals that carried the *R26R^Tomato^*lineage reporter allele revealed an unexpected and significant population of cells in the mutant islets that were *R26R^Tomato^* lineage-positive but lacked insulin expression (Figure 4C-H). Altogether we identified three populations of beta cells in the mutant islets: (i) Insulin-expressing/*R26R^Tomato^*-expressing beta cells, (ii) Insulin-deficient/*R26R^Tomato^*-expressing beta cells, and (iii) Insulin-expressing beta cells that lacked *R26R^Tomato^*lineage label. By comparison, nearly all beta cells in *Ins1-cre* control islets co-expressed insulin and the *R26R^Tomato^*lineage label (Figure 4I). These data suggest that genetic deletion of *Dhps* in the beta cells, and thus loss of eIF5A^HYP^, results in altered synthesis of proteins required for beta cell identity, which may in turn disrupt beta cell function.

As further confirmation that DHPS/eIF5A^HYP^ is required for beta cell identity and function, we examined beta cells from DHPS^ΔBETA^ and *Ins1-cre* mice at 6 weeks-of-age for known markers of identity, namely Slc2a2 (Glut2), Pdx1, and MafA. Immunofluorescence analysis showed punctate, intracellular expression of Glut2 in DHPS^ΔBETA^ lineage-labeled beta cells compared with membrane-bound expression of Glut2 in *Ins1-cre* lineage-labeled beta cells (Figure 5A-F). Furthermore, quantitative image analysis determined that the lineage-labeled (Tomato-expressing), insulin-deficient beta cells in the DHPS^ΔBETA^ mutants were deficient in Glut2 (Figure 5G). We performed a similar analysis for expression and localization of the transcription factors Pdx1 and MafA and observed the lineage-labeled (Tomato-expressing), insulin-deficient beta cells in the DHPS^ΔBETA^ mutants showed little or no Pdx1 or MafA (Figure 5 H-U). Cells in the DHPS^ΔBETA^ mutant islets that maintained expression of Pdx1 likely represent the delta cell population (Figure 5K). Taken together, our data demonstrate that the synthesis of critical proteins for beta cell identity and function is facilitated by DHPS/eIF5A^HYP^, suggesting that beta cell maturation is regulated at the level of mRNA translation.

**Figure 5.**
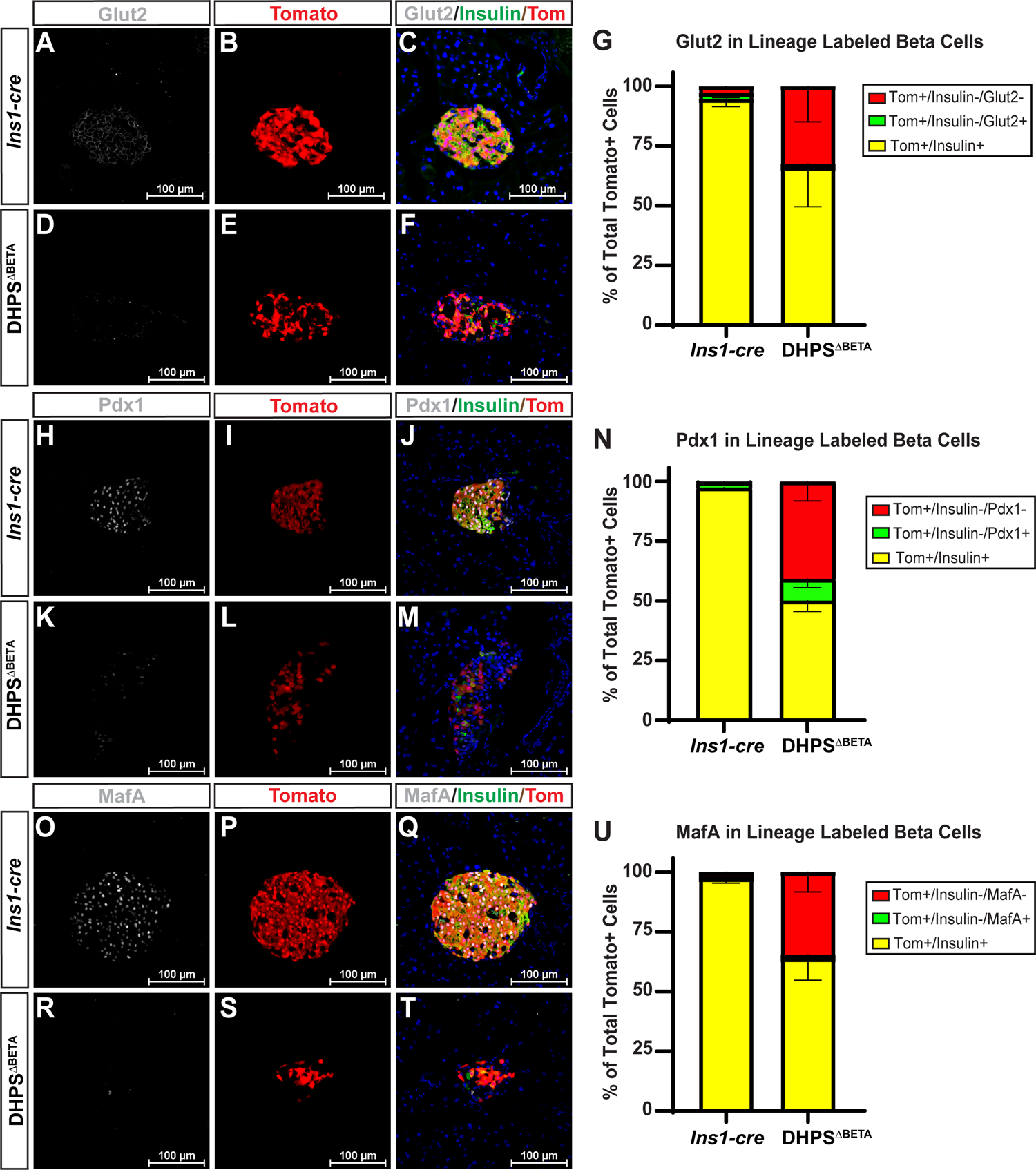
Markers of beta cell identity and function are reduced in DHPS islets. Immunofluorescence on pancreas tissue from 6-week-old (A – C) *Ins1-cre* and (D – F) DHPS^ΔBETA^ mice to determine co-expression of insulin (green), *R26R^Tomato^* beta cell lineage reporter (Tomato, red) and Glut2 (visualized in white). (G) Quantification of lineage-labeled cells with or without Glut2 expression as a percentage of total tomato+ cells. (H - M) Immunofluorescence for Pdx1 and (N) quantification of lineage-labeled cells with or without Pdx1 expression as a percentage of total tomato+ cells. (O-T) Immunofluorescence for MafA and (U) quantification of lineage-labeled cells with or without MafA expression as a percentage of total tomato+ cells. Sytox was used to identify nuclei (blue). Data are represented as mean +/- SEM. (n = 3/group)

## DISCUSSION

Identifying mechanisms that drive beta cell dysfunction is paramount to developing therapeutics for diabetes. The treatment of diabetes currently involves the use of exogenous insulin and pharmacological intervention; however, beta cell replacement or regeneration would represent a step toward the restoration of beta cell mass and function. The development of such cellular therapeutics requires a detailed understanding of the mechanisms controlling the development of the beta cell throughout all stages of its life. In this study, we have identified that there exists a translational regulatory mechanism in the beta cell, driven by DHPS/eIF5A^HYP^, that facilitates the maintenance of beta cell maturation. Specifically, we have shown that beta cell-specific deletion of *Dhps* reduces the synthesis of proteins critical to beta cell identity and function at the stage of beta cell maturation. We have also determined that when this translational regulatory mechanism is altered, there is a rapid and reproducible onset of diabetes.

To fully understand how beta cells develop, these cells must be studied at all stages of significant development in mice – from differentiation in the embryo, to expansion during the first week of postnatal life, to functional maturation around weaning. In our previous work, we characterized a pancreas-specific deletion of *Dhps* (DHPS^ΔPANC^), which showed exocrine insufficiency but no change in beta cell differentiation or abundance in the embryo (26). In the current study, we observed proper perinatal beta cell mass in the DHPS^ΔBETA^ mice but reduced mass and function after weaning. Therefore, our data demonstrate that embryonic and perinatal beta cells do not require the specialized translational regulation provided by DHPS/eIF5A^HYP^ in early stages of beta cell differentiation and expansion. However, under circumstances of cellular stress, such as a rapid change in nutrition or increased metabolic load, eIF5A^HYP^ facilitates the critically important increased rate of protein synthesis. In particular, we showed that in the absence of DHPS, eIF5A is not activated via hypusination, resulting in an inability to increase the synthesis of specific proteins including GLUT2, MAFA, PDX1, UCN3, and INS, which are needed for beta cells to mature and respond properly to new metabolic cues. It is clear from our previous (26) and current work that eIF5A^HYP^ is needed for the translation of only a subset of transcripts in certain cell types and at specific times, with the greatest induction of eIF5A^HYP^ being in response to stress.

In mice, beta cell maturation occurs around weaning age, when the animals shift from a fat-rich milk diet to that of a carbohydrate-rich diet (12). Resultantly, the beta cells must rapidly acquire the machinery to respond to new metabolic cues, which is a form of cellular stress. We propose that the beta cell has developed a translational regulatory mechanism that employs DHPS/eIF5A^HYP^ to increase the rate at which proteins critical for the beta cell secretory function must be synthesized to permit the beta cell to respond appropriately to new metabolic cues. Our studies also extend published work showing that loss of *Dhps* in the adult beta cell in the setting of high fat diet, another cellular stress, results in an inability for beta cells to undergo compensatory proliferation (28). If we consider altogether the work from our group in the exocrine pancreas (26,27) and beta cells (this study) as well as work from others who have studied the brain (41), we propose that eIF5A^HYP^ is a gatekeeper of specialized translation that permits professional secretory cells (exocrine, endocrine, neuronal) to maintain their functional capacity during times of stress. In fact, perhaps it is all metabolically responsive secretory cells that require a special translation factor to facilitate the integrity of specific protein production during times of induced/increased demand.

Studies in yeast support this idea that eIF5A^HYP^ is required to increase the rate of read-through translation elongation for certain transcripts, and that losing the activation of this specialized translation factor reduces the rate of protein synthesis but does not halt global translation (21). In this simplest eukaryotic organism, challenging sequences encoding stretches of consecutive amino acids such as proline require eIF5A^HYP^ for efficient translation elongation (20,24). Further investigation is required to determine if there are sequences within certain beta cell genes that specifically draw the function of eIF5A^HYP^. That said, our work provides an interesting perspective that perhaps driving an increase in translation in the setting of beta cell dysfunction could overcome or possibly reverse the beta cell defects that contribute to early dysfunction and the progression to diabetes.

In short, our mouse model has revealed a regulatory mechanism driven by DHPS/eIF5A^HYP^ that is essential for metabolic health and beta cell function. Furthermore, this mouse model could prove even more useful in the search for understanding pre-diabetes. It is well documented that beta cell dedifferentiation and dysfunction are hallmarks of diabetes pathogenesis (42,43); however the events that precede these changes are difficult to study without biomarkers that identify the stages before the measurable clinical symptoms. The predictable and reproducible progression of loss of beta cell maturation, induced hyperglycemia, and diabetes onset in our DHPS^ΔBETA^ mutants make this a model that can be exploited to identify biomarkers of pre-diabetes. This must be explored.

## ACKNOWLEDGEMENTS

This work was supported by funding to TLM from the Juvenile Diabetes Research Foundation (JDRF) (5-CDA-2016-194-A-N) and the National Institutes of Health (NIH) (R01DK121987-01A1), and funding to SRR from the NIH–IOTN: Data Management and Resource-Sharing Center (U24CA232979). The mass spectrometry work performed by the Indiana University School of Medicine Center for Proteome Analysis. Acquisition of the IUSM Proteomics instrumentation was provided by the Indiana University Precision Health Initiative. The proteomics work was supported, in part, by the Indiana Clinical and Translational Sciences Institute funded in part by funds (UL1TR002529) from the NIH, National Center for Advancing Translational Sciences, Clinical and Translational Sciences Award, and the Cancer Center Support Grant for the IU Simon Comprehensive Cancer Center (P30CA082709) from the National Cancer Institute. Teresa L. Mastracci is the guarantor of this manuscript.

## AUTHOR CONTRIBUTIONS

Author contributions are as follows: TLM designed the study; CTC, EKAB, SR, CBPV, CDR, PJC, LRP, MAR, and TLM performed the research; CTC, EKAB, SR, CBPV, CDR, LRP, MAR, and TLM analyzed data; CTC, EKAB and TLM wrote the paper. All authors edited and approved the final draft of the manuscript.

