## Supplemental Figures for "Deoxyhypusine synthase is required for the translational regulation of pancreatic beta cell maturation"

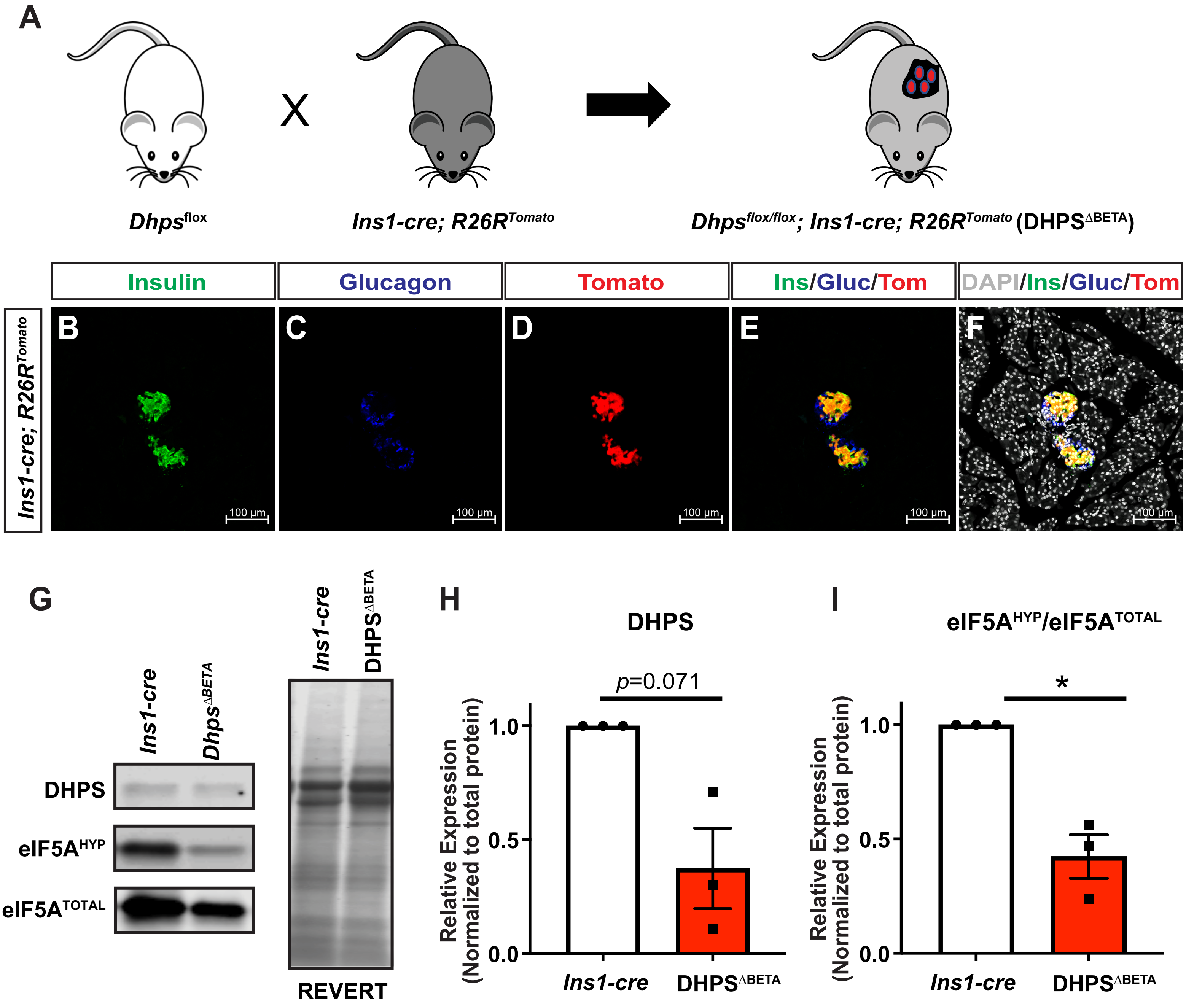


**Supplemental Figure 1. A mouse model of beta cell-specific genetic deletion of *Dhps*.**
(A) Mouse alleles used to generate the beta cell-specific deletion of *Dhps* ($DHPS^{\Delta BETA}$), which includes a reporter allele to permit the lineage labeling of beta cells. Pancreas tissue from control animals (*Ins1-cre;R26R*$^{Tomato}$) was evaluated by immunofluorescence for expression of (B) insulin, (C) glucagon, and (D) tomato to demonstrate specific cre activity in the beta cells. (E) Islets showing co-expression of Tomato in the insulin-expressing beta cells. (F) DAPI, a nuclear marker, was added to define all cells. (G) Western blot analysis of islets from 4 week-old $DHPS^{\Delta BETA}$ mutants for expression of DHPS, $eIF5A^{HYP}$, $eIF5A^{TOTAL}$ and total protein (as visualized by REVERT™). (H) Densitometric data for DHPS and the ratio of $eIF5A^{HYP}/eIF5A^{TOTAL}$ are represented as mean +/- SEM, n = 3, * $p < 0.05$.

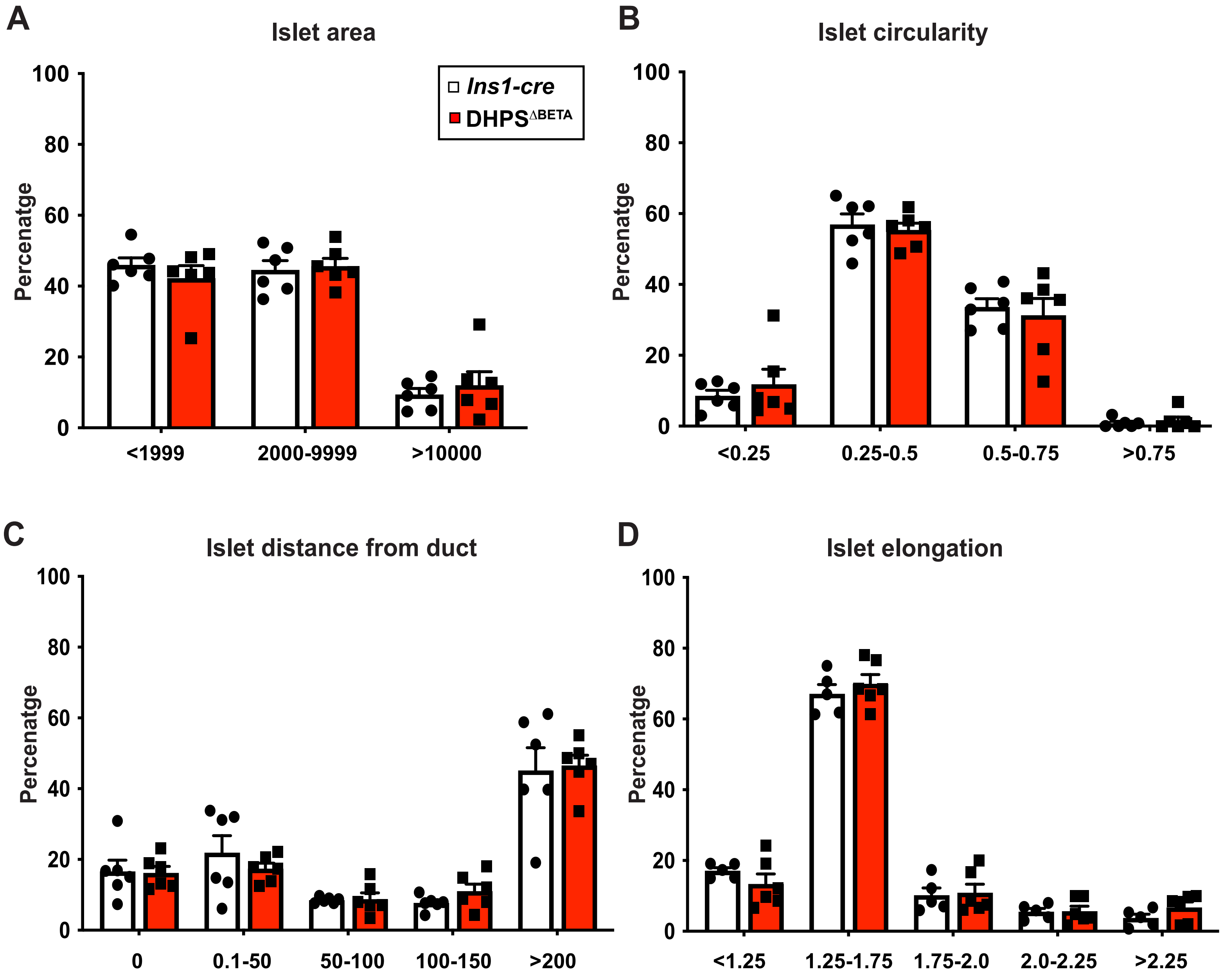


**Supplemental Figure 2. Perinatal islet morphology in the absence of *Dhps*.**
(A) Pancreas tissue from 1 week-old *Ins1-cre* controls and DHPS$^{\Delta BETA}$ mutant mice was evaluated for islet size, which was categorized as small (<1999 μm), medium (2000-9999 μm) and large (>10000 μm). (B) Islet circularity was measured on a scale of 0 - 1, where 1 is a perfect circle. (C) Islet to duct distance was measured in μm, with 0 representing an islet that is directly adjacent to its closest duct. (D) Islet shape (elongation) was determined by measuring the minimum (shortest) and maximum (widest) Ferret's diameter. All data are reported as percentages; n = 6/group; graphs are displayed as mean +/- SEM.

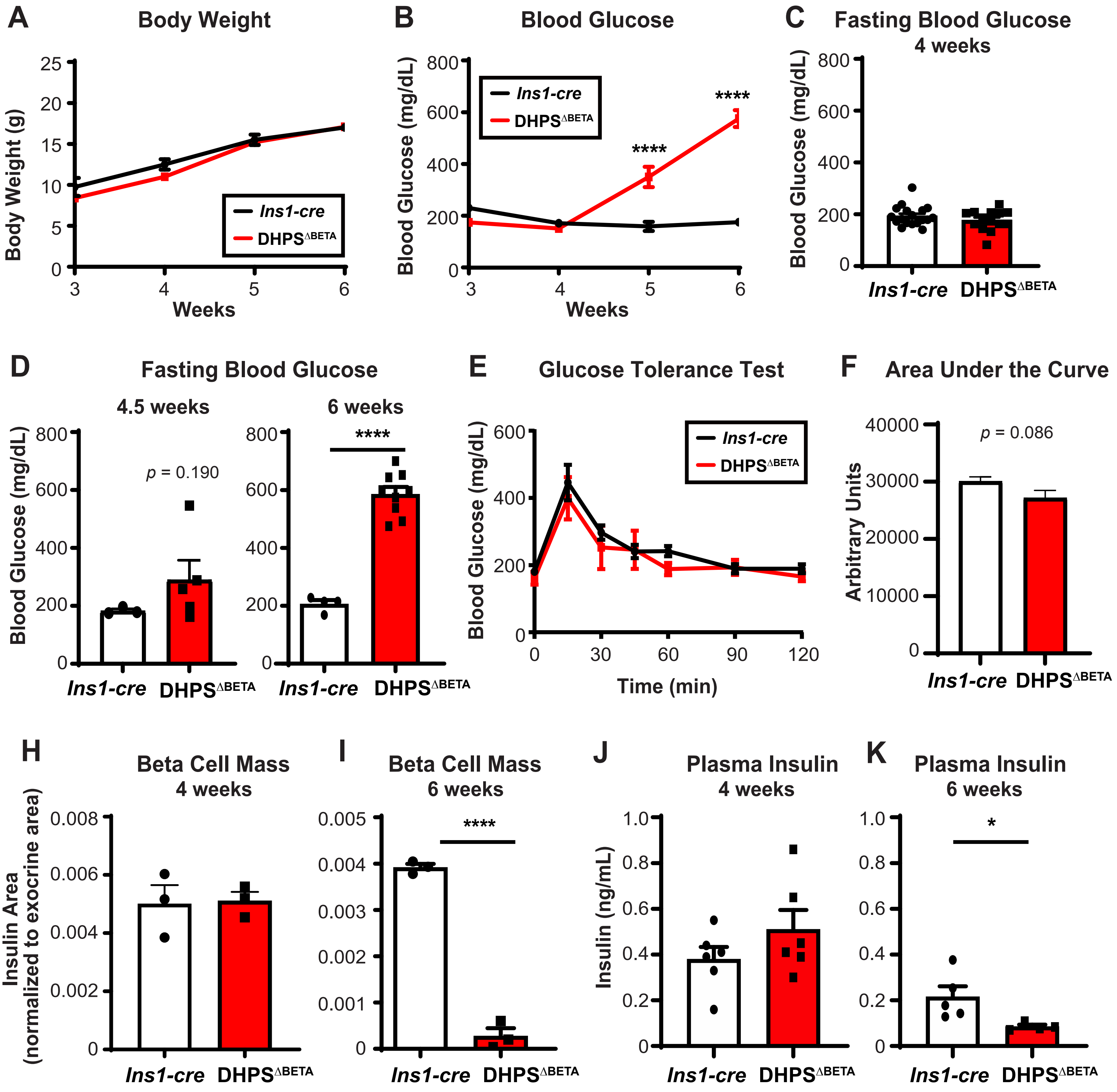


**Supplemental Figure 3. Metabolic and beta cell phenotype of DHPS$^{\Delta BETA}$ female mice**
(A) Body weight and (B) *ad libidum* fed blood glucose were measured in *Ins1-cre* and DHPS$^{\Delta BETA}$ mice weekly beginning at weaning (3 weeks-of-age)(n = 4 - 10/group). Five hour fasting blood glucose was measured in control and DHPS$^{\Delta BETA}$ mice at (C) 4 weeks-of-age, (D) 4.5 weeks-of-age, and 6 weeks-of-age (n = 3 -18/group). (E) Glucose Tolerance Tests (GTT) were performed on 4 week-old mice (n = 7/group). (F) Area under the curve was quantified from the GTT data. From immunohistochemical staining, beta cell mass was quantified as insulin area normalized to pancreas area in (H) 4 week-old and (I) 6 week-old *Ins1-cre* controls and DHPS$^{\Delta BETA}$ mutants (n = 3/group). Insulin levels were measured from plasma collected from (J) 4 week-old and (K) 6 week-old *Ins1-cre* controls and DHPS$^{\Delta BETA}$ mutants (n = 4-6/group). Quantitative data are represented as mean +/- SEM. * $p<0.05$; *** $p<0.001$; **** $p<0.0001$.

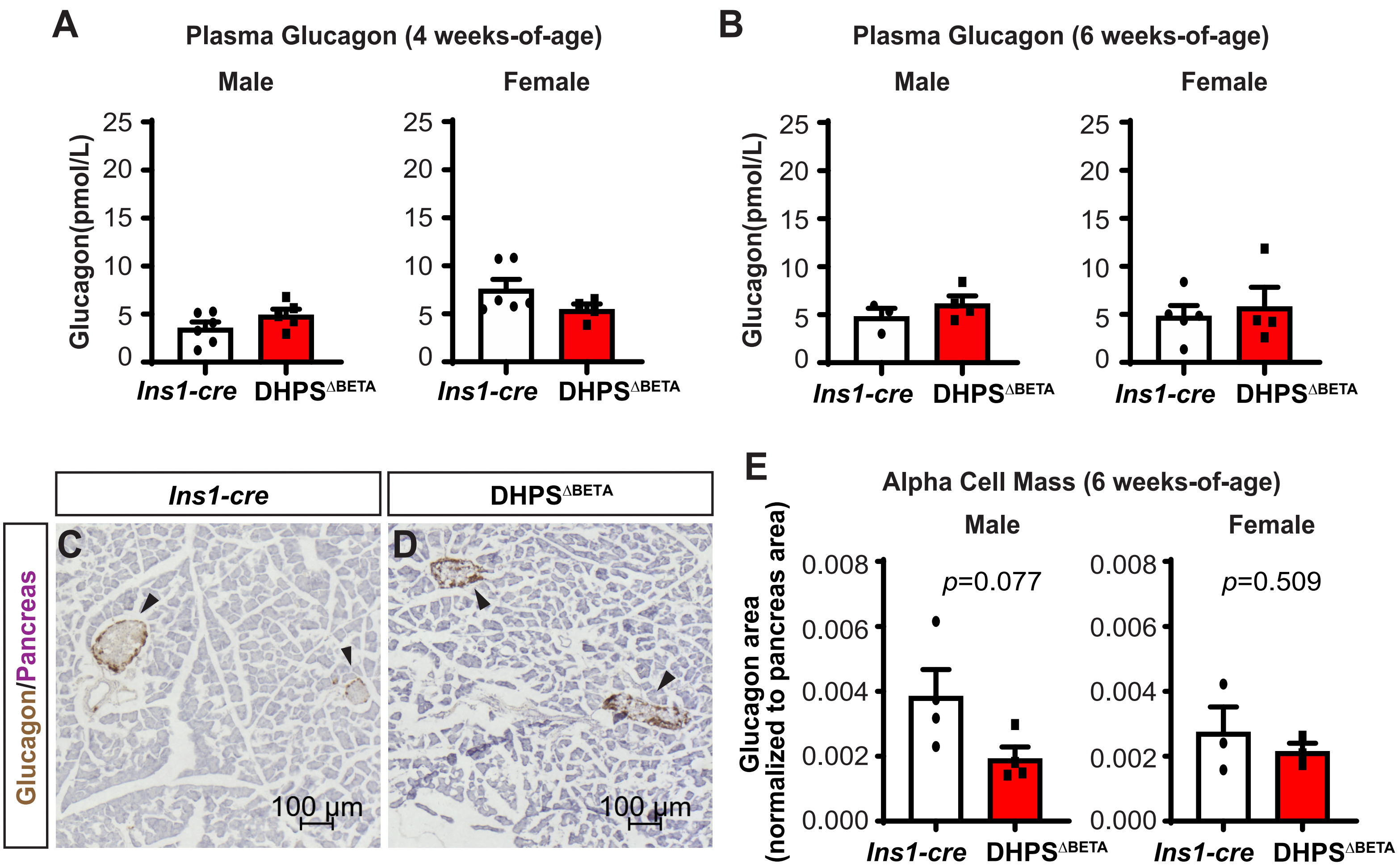


**Supplemental Figure 4. Beta cell-specific loss of *Dhps* does not impact alpha cell growth and function.** Glucagon levels were measured from plasma collected from (A) 4 week-old and (B) 6 week-old male and female *Ins1-cre* and DHPS$^{\Delta BETA}$ mice (n = 3 - 6/group). Immunohistochemistry for Glucagon (brown) was performed on pancreas tissue from 6 week-old (C) *Ins1-cre* control and (D) DHPS$^{\Delta BETA}$ mutant male and female mice (n = 3 - 4/group). All sections were counterstained with Eosin (purple) to define pancreas tissue. (E) Alpha cell mass was quantified as glucagon area normalized to pancreas area. Data are represented as mean +/- SEM.

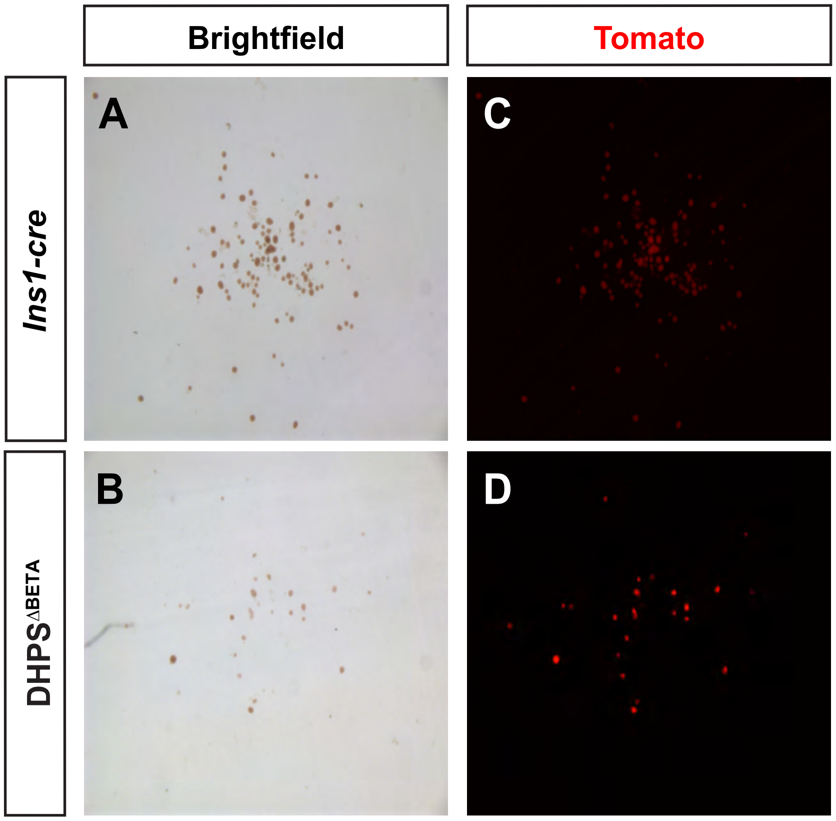


**Supplemental Figure 5. Wholemount images of islet isolations from *Ins1-cre* and DHPS$^{\Delta BETA}$ mice**
Brightfield images of islets isolated from 4 weeks-old (A) *Ins1-cre* and (B) DHPS$^{\Delta BETA}$ mice. Islet preparations were processed to remove any remaining exocrine cell contamination after collagenase digestion and separation by ficol gradient. Islets from the (C) *Ins1-cre* and (D) DHPS$^{\Delta BETA}$ mice were also imaged to visualize the expression of the *R26R*$^{Tomato}$ reporter (beta cell lineage label; red).
