## Supplemental Methods for "Deoxyhypusine synthase is required for the translational regulation of pancreatic beta cell maturation"

**SUPPLEMENTAL MATERIALS AND METHODS**

**Table S1.** ***Antibodies and Dyes for Immunofluorescence, Immunohistochemistry and Western blot***

| Procedure | Antibody | Company | Catalog number | Dilution |
|---|---|---|---|---|
| IF, IHC | guinea pig anti-Insulin | Agilent | A056401-2 | 1:500 |
| IF, IHC | rabbit anti-Glucagon | Cell Signaling | 2760S | 1:500 |
| IF | rabbit anti-Glut2 | Abcam | ab54460 | 1:200 |
| IF | rabbit anti-Pdx1 | Millipore Sigma | 07696 | 1:500 |
| IF | rabbit anti-MafA | Cell Signaling | 79737S | 1:500 |
| IF | goat anti-CarboxypeptidaseA | R & D Systems | AF2765 | 1:500 |
| Western | mouse anti-DHPS | Santa Cruz | sc-365077 | 1:2000 |
| Western | rabbit anti-$eIF5A^{HYP}$ | Millipore | ABS1064-I-25UL | 1:2000 |
| Western | mouse anti-eIF5A | BD Biosciences | BDB611977 Clone 26 | 1:2000 |
| Western | IRDye 680RD | LI-COR Biosciences | 926-68072 | 1:10000 |
| Western | IRDye 800CW | LI-COR Biosciences | 926-32213 | 1:10000 |
| IF | DyLight™ 405 | Jackson ImmunoResearch | 706-475-148 | 1:500 |
| IF | Alexa 488 | Jackson ImmunoResearch | 705-545-147 | 1:500 |
| IF | Alexa 647 | Jackson ImmunoResearch | 711-605-152 | 1:500 |
| IHC | horseradish peroxidase (HRP)-conjugated donkey anti-guinea pig | Jackson ImmunoResearch | 706-035-148 | 1:500 |
| IHC | HRP-conjugated donkey anti-rabbit | Jackson ImmunoResearch | 711-035-152 | 1:500 |
| IF | DAPI | Sigma | 10236276001 | 1:1000 |
| IF | Sytox | Fisher | S7020 | 1:5000 |

***Pancreatic Islet Isolation***

Pancreata of $DHPS^{DBETA}$ and *Ins1-cre* mice were injected with 0.7 mg/mL of collagenase (Sigma) in cold Hank's Balanced Salt Solution (HBSS; Fisher) containing 0.3% bovine serum albumin (Fisher) into the common bile duct. The pancreas, inflated with the collagenase solution, was then dissected and incubated in a 37$^{o}$C water bath for 16 minutes. The digested pancreas was placed on ice, 30 mL of HBSS/BSA was added and the tissue centrifuged at 1000 rpm for 1 minute. Following removal of the supernatant, the pellet was dissociated in 10 mL of fresh HBSS/BSA using a 14 gauge bore pipet attached to a 30 mL syringe (Fisher). The disrupted tissue is then passed through a strainer, rinsed with HBSS/BSA and pelleted by centrifugation for 2 minutes at 1280 rpm. The pelleted tissue was resuspended in 10 mL of Histopaque 1100 (Fisher) and 10 mL of HBSS/BSA was carefully layered over the Histopaque to create two distinct layers. The gradient was then centrifuged for 18 minutes at 2080 rpm. The top layer containing the islets was removed and filtered to collect the isolated islets. Islets were rinsed from the filter using RPMI 1640 culture media (Gibco/ThermoFisher) containing 10% fetal bovine serum (Fisher) and collected into a non-tissue culture treated 6-well dish (Fisher). Islets were then handpicked, counted and collected into 1.5 mL tube (Eppendorf) and centrifuged at 4$^{o}$C for 5 minutes at 10,000 rpm. Any remaining culture media was removed and the pelleted islets and used for either western blot or quantitative mass spectrometry analysis.

***Quantitative mass spectrometry***

Islets were lysed in in 8 M urea, 50 mM Tris.HCl pH 8.5 (25 µL), and then sonicated in a Bioruptor® sonication system (Diagende Inc.) with 30 sec/30 sec on/off cycles for 15 minutes in a water bath at 4$^{o}$C. Protein concentrations were determined using a Bradford protein assay (BioRad) employing vendor provided protocols. Protein samples in equal amounts (15 ug) were reduced with 5 mM tris(2-carboxyethyl)phosphine hydrochloride for 30 min at RT and then free Cysteines were alkylated with 10 mM chloroacetamide for 30 min at RT in the dark. Samples were diluted with 100 mM Tris-HCl, pH 8.5 to a final urea concentration of 2 M and digested overnight with Trypsin/Lys-C Mix Mass Spectrometry (Promega Corporation , V5072, 1:100 protease:substrate ratio)(36–38).

Peptides were desalted on 50 mg Sep-Pak® Vac (Waters Corporation Milford) employing a vacuum manifold (Waters Corporation Milford). After elution from the column in 70% ACN, 0.1% FA, peptides were dried by speed vacuum and resuspended in 24 µL of 50 mM triethylammonium bicarbonate. Peptide concentration was measured using Pierce Quantitative Colorimetric Peptide Assay Kit (Thermo Fisher Scientific) to ensure that an equal amount of each was labeled. Samples were then Tandem Mass Tag (TMT) labeled with 0.2 mg of reagent for two hours at room temperature (Thermo Fisher Scientific, 90110). Labelling reactions were quenched with hydroxylamine at room temperature 15 minutes. Labelled peptides were then mixed and dried by speed vacuum.

Each peptide mixture was resuspended in 0.1% TFA (trifluoroacetic acid) and fractionated on Pierce™ High pH reversed-phase peptide fractionation spin columns using vendor methodology (Thermo Fisher Scientific, 84868). Each fraction was dried by speed vacuum and resuspended in 24 uL 0.1% formic acid (FA).

Nano-LC-MS/MS analyses were performed on an EASY-nLC™ HPLC system (Thermo Scientific) coupled to an Orbitrap Fusion™ Lumos™ mass spectrometer (Thermo Fisher Scientific). One third of each fraction was loaded onto a reversed phase PepMap™ RSLC C18 column (2 µm, 100 Å, 75 µm x 50 cm) with Easy-Spray tip at 400 nL/min. Peptides were eluted from 4-28% B over 160 minutes, 28%-35% B over 5 mins, 35-50% B for 14 minutes, and dropping from 50-10%B over the final 1 min (Mobile phases A: 0.1% FA, water; B: 0.1% FA, 80% Acetonitrile). Mass spectrometer settings include capillary temperature of 275 $^{o}$C and ion spray voltage was kept at 2.5 kV. The mass spectrometer method was operated in positive ion with a 4 sec cycle time data-dependent acquisition method with advanced peak determination and Easy-IC (internal calibrant). Precursor scans (m/z 400-1750) were done with an orbitrap resolution of 120000, RF lens% 30, maximum inject time 50 ms, standard AGC target, including charges of 2 to 6 for fragmentation with 60 s dynamic exclusion.

The following settings were used for HCD $MS^2$ scans: Isolation mode = Quadrupole; Isolation Offset = Off; Isolation Window = 1; Multi-notch Isolation = False; Scan Range Mode = Auto Normal; First Mass = 100; Activation Type = HCD; Collision Energy Mode = Fixed;

Collision Energy (%) = 35; Detector Type = Orbitrap; Orbitrap Resolution = 50k; Data type = Centroid; scan range define first mass 100; AGC target normalized 20%; maximum IT mode: dynamic. The data were recorded using Thermo Scientific Xcalibur (4.3) software (Thermo Fisher Scientific).
